# Melatonin: The smart molecule that differentially modulates autophagy in tumor and normal placental cells

**DOI:** 10.1101/385948

**Authors:** Lucas Sagrillo-Fagundes, Josianne Bienvenue-Pariseault, Cathy Vaillancourt

**Affiliations:** INRS-Institut Armand-Frappier and BioMed Research Centre, Laval, QC, Canada and Center for Interdisciplinary Research on Well-Being, Health, Society and Environment, Université du Québec à Montréal, Montréal, QC, Canada.

**Keywords:** BeWo cells, villous cytotrophoblast, pregnancy, melatonin, autophagy, Nrf2

## Abstract

**Highlights:**

- H/R induces autophagy and Nrf2 in tumoral and primary trophoblast cells.
- Melatonin inhibits autophagy and Nrf2 under H/R, inducing BeWo cell death.
- Melatonin increases autophagy and Nrf2 under H/R conditions, promoting primary villous trophoblast cells survival.

**Abstract:** Hypoxia/reoxygenation (H/R) induces oxidative damage and apoptosis. These consequences activate autophagy, which degrades damaged cellular content, as well as inducing the nuclear factor (erythroid-derived 2)-like 2 (Nrf2) transcription factor, and thereby the expression of protective genes. Melatonin has protective roles in normal cells and cytotoxic actions in cancer cells, with effects involving autophagy and Nrf2 pathways. The current study shows melatonin to differentially modulate autophagy and Nrf2 pathways in tumor and normal placental cells exposed to H/R. BeWo, a human placental choriocarcinoma cell line, and primary villous cytotrophoblasts isolated from normal term placenta, were maintained in normoxia (8% O_2_) for 24 h or exposed to hypoxia (0.5% of O_2_ for 4 h) followed by 20 h of normoxia, creating a H/R, in the presence or absence of 1 mM melatonin. Melatonin induced a 7-fold increase in the activation of 5’ adenosine monophosphate-activated protein kinase (AMPK)α, an upstream modulator of autophagy, rising to a 16-fold increase in cells co-exposed to H/R and melatonin, compared to controls. H/R induced autophagosome formation via the increased expression of Beclin-1 (by 94 %) and ATG7 (by 97%). H/R also induced autophagic activity, indicated by the by the 630% increase in P62, and increased Nrf2 by 314%. In H/R conditions, melatonin reduced autophagy by 74% and Nrf2 expression by 66%, leading to BeWo cell apoptosis. In contrast, in human primary villous cytotrophoblasts, H/R induced autophagy and Nrf2, which melatonin further potentiated, thereby affording protection against H/R. This study demonstrates that melatonin differentially modulates autophagy and the Nrf2 pathway in normal vs. tumor trophoblast cells, being cytoprotective in normal cells whilst increasing apoptosis in tumoral trophoblast cells.

## Introduction

Macroautophagy, herein referred to as autophagy, is a highly conserved detoxifying mechanism involving the catabolism of damaged proteins and organelles [1]. Autophagy shows low levels of activity under basal conditions, being inhibited by the cellular sensor, the mechanistic target of rapamycin (mTOR). However, autophagy is activated in suboptimal conditions, such as hypoxia/reoxygenation (H//R) or amino acid starvation (reviewed in [2]). Beclin-1 is an important initiator of autophagy via its activation of the ATG (autophagy-related) proteins. ATG proteins build a double-membrane vesicle, autophagosome, which engulfs cargo to be degraded in lysosomes. The consequent release of simpler structures can restore cellular energy levels and inhibit the deleterious effects of reactive species of oxygen (ROS) [3, 4]. Autophagy upregulates the transcription factor nuclear factor (erythroid-derived 2)-like 2 (Nrf2, also called NFE2L2), by the autophagy carrier sequestosome-1/P62 (SQSTM1/P62) [5]. Nrf2 induces defenses against oxidative and other stressors, including by binding to the consensus antioxidant response element (ARE) in their promoters. As with autophagy, Nrf2 is activated in during hypoxia in both normal and cancer cells, including placental cells [6–8].

Alterations in oxygenation are common, reducing cell viability including by increasing ROS and oxidative stress, thereby leading to oxidation and damage of proteins, DNA and lipids [9, 10]. Under such challenge, autophagy is activated leading to increased catabolism of damaged cellular components. BeWo cells, a placental choriocarcinoma model, are frequently utilized to investigate placental physiology, given their ability to synthesize human chorionic gonadotropin (hCG) and their ability to mimic the differentiation of villous cytotrophoblasts (vCTB) into syncytiotrophoblast (STB) [11, 12]. During altered oxygenation, both BeWo and primary trophoblast cells show increased ROS and cell death, thereby inducing autophagic activity, which is modulated by the 5’ adenosine monophosphate-activated protein kinase (AMPK)α and the protein phosphatase 2c (PP2Ac), cellular sensors that are activated to enhance cell survival [13–16].

Melatonin is produced by most cell types, across different tissues and organs. Melatonin is a strong antioxidant, anti-inflammatory and optimizer of mitochondria functioning in non-tumor cells [17, 18]. In contrast, melatonin is cytotoxic in tumor cells, where it has pro-apoptotic and antiproliferative effects [19]. In human placental trophoblastic cells, we have previously shown melatonin to reverse H/R-induced elevations in oxidative stress and cell death, mediated via melatonin effects on inflammation and autophagy [20]. In human choriocarcinoma cells, melatonin disrupts the permeability of the mitochondrial membrane, leading to intrinsic apoptosis [21]. The mechanisms underlying these distinctive effects of melatonin normal vs tumoral placental cells have still to be determined. The comparative effects of melatonin on autophagy and Nrf2 levels in normal vs tumoral placental cells have yet to be investigated. The current study shows that under H/R conditions, the autophagic activity and related pathways are increased in BeWo cells, acting to protect these cells against apoptosis. Melatonin treatment blocks the rise in autophagy in BeWo cells, thereby contributing to their apoptosis. In primary cells, H/R also enhances autophagic activity, which is further increased by melatonin, thereby contributing to cell survival.

## Materials and methods

### Cell culture

BeWo cells (CCL-98 clone), from American Type Culture Collection (ATCC; Rockville, MD), were cultured in Dulbecco’s Modified Eagle Medium (DMEM)/F-12 without phenol red and supplemented with 10% fetal bovine serum (FBS; Hyclone, Tempe, AZ). Human primary vCTB, were obtained from term placentas arising from the spontaneous vaginal delivery of uncomplicated pregnancies after the ethical approval from the CHUM-St-Luc Hospital (Montreal, QC, Canada) and informed patient consent. vCTB cells were isolated based on the classic method trypsin-DNase/Percoll [22], modified by our group [23]. Briefly, placenta was exposed to four consecutive digestions with trypsin and DNAse. The supernatant of each bath was collected and pooled. The pooled pellets were added to a Percoll gradient, centrifuged, and then the layers containing 40 - 50% of density, which contained mainly villous trophoblast, were collected. Following the isolation, mononuclear villous trophoblasts were immunopurified using the autoMACS™ (Myltenyi Biotec, Santa Barbara), as described previously [24, 25]. All vCTB preparations used in this study were at least of 95% purity after cell sorting. vCTB were cultured in DMEM–High glucose, containing 10% FBS and 1% penicillin-streptomycin (Hyclone).

Cells were plated at the densities of 2.5 × 10^6^ cells/ml (BeWo cells) and 3.0 × 10^6^ cells/ml (primary vCTB) and promptly put under normoxia (8% O_2_, 5% CO_2_, 87% N_2_) in Modular Incubator Chambers (Billups-Rothenberg, San Diego, CA, USA) for 6 h in order to ensure cell adherence, as previously described. Cells were washed thoroughly with warm media and then maintained under normoxia (8% O_2_,) for 24 h or exposed to hypoxic conditions (0.5% O_2_,) for 4 h followed by 20 h of normoxia, creating a hypoxia/reoxygenation effect (**Fig. 1a)**. Both BeWo and vCTB cells were treated with 1mM melatonin (Sigma-Aldrich, Oakville, ON, Canada) for 24 h or with its vehicle control, 0.1% dimethyl sulfoxide (DMSO; Sigma-Aldrich). Cells were treated for 24 h with Rapamycin ((300 nM) Tocris, Burlington, ON, Canada) and 3-Methyladenine ((5 mM) 3-MA, Tocris), as inducer and inhibitor of autophagy, respectively. Bafilomycin A1 ((10nM) Tocris) was added to cells during the last 2 h of culture to inhibit the fusion of the autophagosome with the lysosome. Sulforaphane ((10μM); Enzo Life Science, New York) was used as an inducer of Nrf2 activity.

### MTT assay

Cell viability was determined by monitoring the conversion of 3-(4,5-dimethylthiazol-2-yl)-2,5-diphenyltetrazolium bromide (MTT) to its insoluble form (formazan), by NAD(P)H-dependent cellular oxidoreductase enzymes. Twenty-four hours after treatment of BeWo cells with DMSO (vehicle control), melatonin 1mM, rapamycin 300 nM or 3-MA 5 mM under normoxia or H/R, 10 µl of MTT, at a final concentration of 5 mg/ml, was added to the media. Four hours after incubation, 100 µl Solubilisation solution (40% (vol/vol) dimethylformamide in 2% (vol/vol) glacial acetic acid) was added to each well to dissolve formazan crystals. After mixing to ensure complete solubilisation, MTT formazan absorbance at 570 nm was measured using with Spectra Max M5 (Molecular Devices). Results are presented as a percent of vehicle control (DMSO).

### hCG secretion

To evaluate the secretion of hCG by BeWo cells exposed to vehicle control (DMSO), melatonin 1 mM, rapamycin 300 nM or 3-MA 5 mM under normoxia or H/R, cell culture media were collected and centrifuged, after which supernatants were stored at −20°C until assayed. The secretion of hCG culture was evaluated by enzyme-linked immunosorbent (ELISA) assay according to manufacturer’s instructions (IBL International; Toronto, ON, Canada). Results are presented as a percent of vehicle control.

### Immunoblotting

To analyze protein expression, BeWo cells and primary vCTB were rinsed with PBS and lysed with ice-cold modified radioimmunoprecipitation (RIPA) buffer (50 mmol/l Tris-HCl pH 7.4, 1% NP-40, 0,25% Na-deoxycholate, 150 mmol/l NaCl and 1 mmol/l EDTA) containing protease and phosphatase inhibitors (Sigma-Aldrich). Protein concentration was determined using the bicinchoninic acid (BCA) protein assay reagent (Pierce Biotechnology, Waltham, MA). Twenty µg of protein were separated on Mini-PROTEAN^®^ TGX™ Precast Protein Gels (4–15% precast polyacrylamide gel) (Bio-Rad, Saint-Laurent, QC, Canada), followed by transfer to Polyvinylidene fluoride (PVDF) membranes (Bio-Rad). Membranes were then incubated with antibodies as described in **Table 1**. Blots were developed with enhanced chemiluminescence reagent (Bio-Rad). Protein levels were expressed as a ratio of a specific band density and total protein stained using Pierce™ Reversible Protein Stain Kit (Pierce Biotechnology), as previously described [26, 27]. Bands were quantified using Image Lab software 6.0 (Bio-Rad).

**Table 1:**
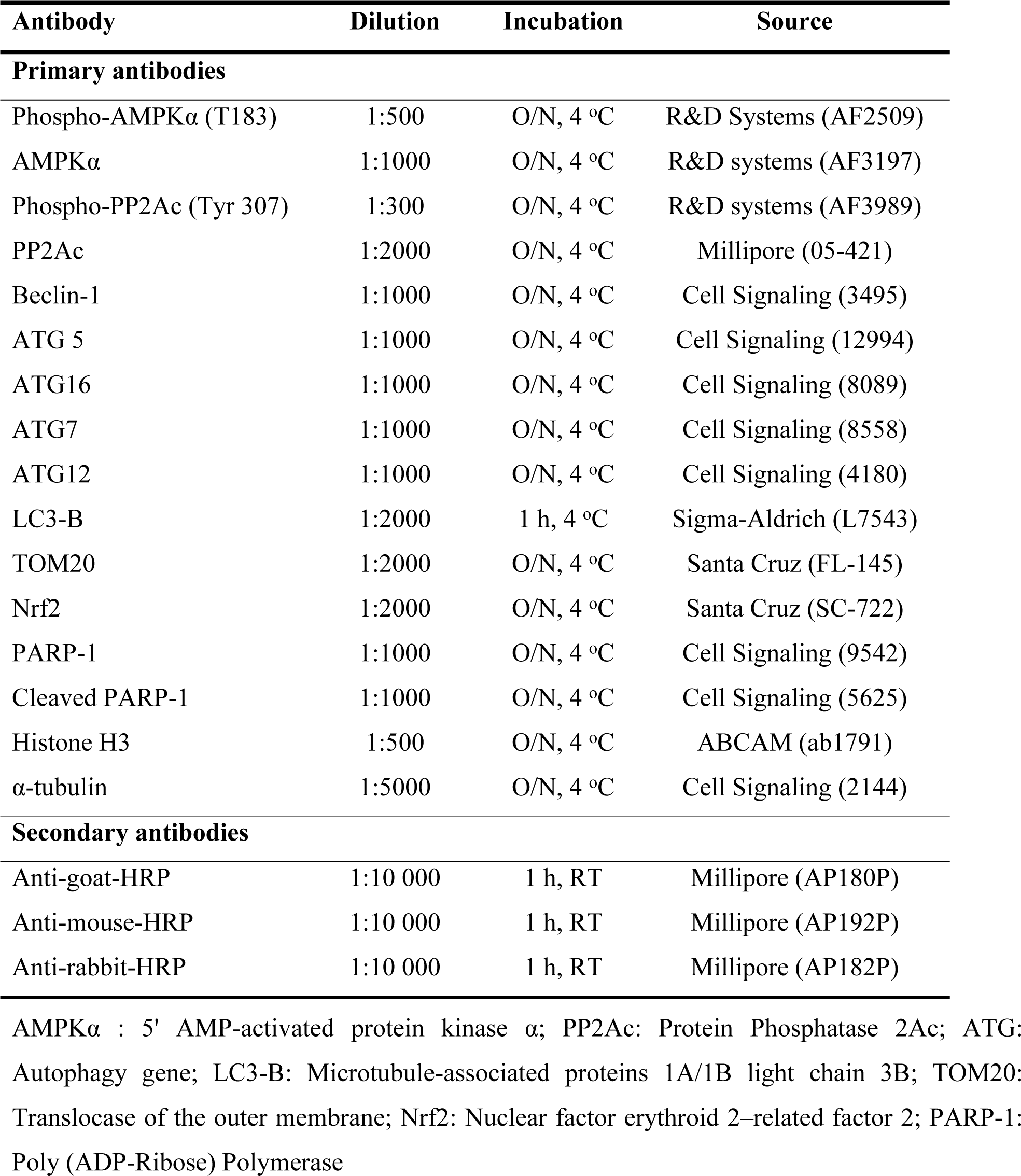
Antibodies used for Western blot analyses.

| Antibody | Dilution | Incubation | Source |
| --- | --- | --- | --- |
| **Primary antibodies** |  |  |  |
| Phospho-AMPKα (T183) | 1:500 | O/N, 4 °C | R&D Systems (AF2509) |
| AMPKα | 1:1000 | O/N, 4 °C | R&D systems (AF3197) |
| Phospho-PP2Ac (Tyr 307) | 1:300 | O/N, 4 °C | R&D systems (AF3989) |
| PP2Ac | 1:2000 | O/N, 4 °C | Millipore (05-421) |
| Beclin-1 | 1:1000 | O/N, 4 °C | Cell Signaling (3495) |
| ATG 5 | 1:1000 | O/N, 4 °C | Cell Signaling (12994) |
| ATG16 | 1:1000 | O/N, 4 °C | Cell Signaling (8089) |
| ATG7 | 1:1000 | O/N, 4 °C | Cell Signaling (8558) |
| ATG12 | 1:1000 | O/N, 4 °C | Cell Signaling (4180) |
| LC3-B | 1:2000 | 1 h, 4 °C | Sigma-Aldrich (L7543) |
| TOM20 | 1:2000 | O/N, 4 °C | Santa Cruz (FL-145) |
| Nrf2 | 1:2000 | O/N, 4 °C | Santa Cruz (SC-722) |
| PARP-1 | 1:1000 | O/N, 4 °C | Cell Signaling (9542) |
| Cleaved PARP-1 | 1:1000 | O/N, 4 °C | Cell Signaling (5625) |
| Histone H3 | 1:500 | O/N, 4 °C | ABCAM (ab1791) |
| α-tubulin | 1:5000 | O/N, 4 °C | Cell Signaling (2144) |
| **Secondary antibodies** |  |  |  |
| Anti-goat-HRP | 1:10 000 | 1 h, RT | Millipore (AP180P) |
| Anti-mouse-HRP | 1:10 000 | 1 h, RT | Millipore (AP192P) |
| Anti-rabbit-HRP | 1:10 000 | 1 h, RT | Millipore (AP182P) |

AMPKα : 5' AMP-activated protein kinase $\alpha$ ; PP2Ac: Protein Phosphatase 2Ac; ATG: Autophagy gene; LC3-B: Microtubule-associated proteins 1A/1B light chain 3B; TOM20: Translocase of the outer membrane; Nrf2: Nuclear factor erythroid 2-related factor 2; PARP-1: Poly (ADP-Ribose) Polymerase

### Immunofluorescence

BeWo cells and primary vCTB were fixed with methanol at – 20 °C for 20 min. Cells were incubated with 2% FBS in PBS for 1 h to eliminate nonspecific antigen binding. BeWo cells were stained overnight (4 °C) for LC3B (1:150) or cytochrome c oxidase iv (cox iv) (1:300). vCTB were stained overnight (4 °C) for LC3-B (1:150) and Nrf2 (1:150). After the incubation with the primary antibodies, both cell types, were then incubated with anti-mouse or anti-rabbit conjugated to Alexa Fluor fluorescent dye. Cell nuclei were stained with 4’,6-diamidino-2-phenylindole (dapi) present in the antifade reagent, prolong™ gold (Thermo Fisher scientific). Immunofluorescence was analyzed at 400 x using a Leica DMRE fluorescence microscope (Ceerfield, il) with a Cooke sensicam high-performance digital CCD camera (Romulus, mi).

### Nucleus isolation

BeWo cells (2.0 × 10^6^ cells) were harvested with TrypLE and had the nuclei isolated following the guidelines of the isolation kit, NE-PER Nuclear and Cytoplasmic Extraction (Thermo Fisher Scientific). The protein yield of the nuclear and the cytosolic extracts was measured with the BCA protein assay. The purity of the nuclear and the cytosolic extracts were validated with the western blotting of the protein Histone H3 (nuclear fraction) and α-tubulin (cytosolic fraction).

### Statistical analysis

All data represent at least four different BeWo cell passages or three primary vCTB cultures. Statistically significant differences (*p* ≤ 0.05) of parametric results were determined by Student’s *t*-test or one-way analysis of variance (ANOVA) and identified by the *post hoc* test of Student-Newman-Keuls. Data were analyzed using GraphPad Prism (version 5.04).

## Results

### Melatonin increases cellular disruption caused by H/R in BeWo cells

As shown in **figure 1b**, BeWo cells exposed to H/R showed a 1.38-fold reduction in viability, compared to normoxia. BeWo cells exposed to H/R and concomitantly with melatonin or 3-MA, an inhibitor of autophagy, also showed lower cell viability (reduction of 1.29 and 1.44-fold, respectively), compared to controls under normoxia. Importantly, in comparison to BeWo cells exposed only to H/R, cells exposed to H/R and melatonin showed significantly lower viability (reduction of 1.19-fold). Cells exposed to H/R and rapamycin, which induces autophagy, showed no differences in cell viability compared to normoxia, suggesting that rapamycin may improve cell viability in BeWo cells exposed to the suboptimal oxygen. In BeWo cells under normoxic conditions, melatonin led to a 1.28-fold reduction in the release of β-hCG, vs DMSO controls (**Fig. 1c**). H/R decreased the release of β-hCG in BeWo cells exposed to vehicle control by 1.38-fold and by 2.1-fold in cells exposed to 3-MA (**Fig. 1c**). In BeWo cells exposed to rapamycin the release of β-hCG was not affected, independently of the levels of oxygenation. The similarity of the viability and β-hCG results in cells treated with 3-MA may suggest that the absence of autophagy affects the global cell viability of BeWo cells exposed to H/R.

**Fig. 1:**
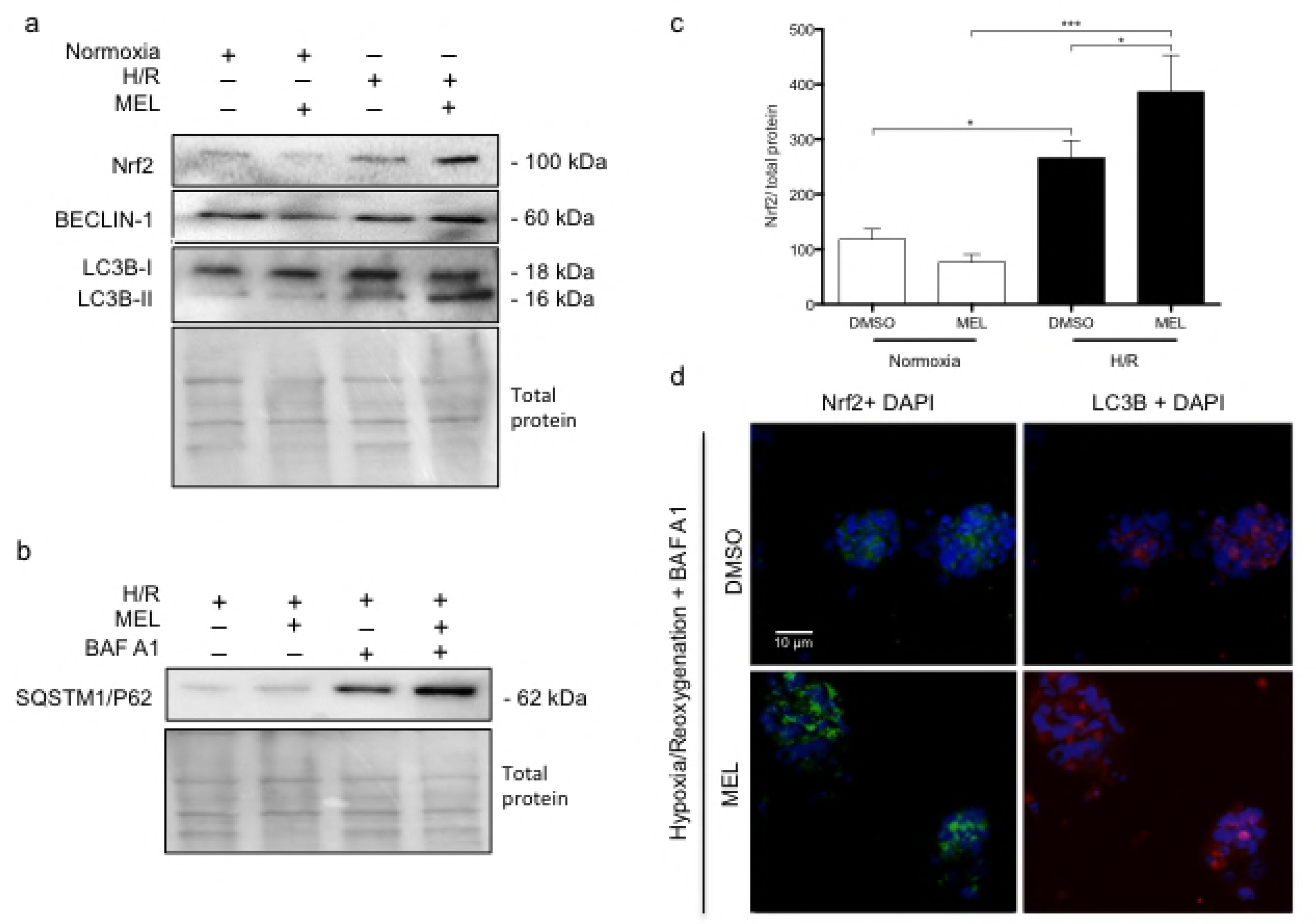
Melatonin decreases the viability of BeWo cells under hypoxia/reoxygenation. (a) BeWo cells were exposed to normoxia (8% O_2_; 5% CO_2_; 87% N_2_) or hypoxia (4 h)/reoxygenation (20 h) (H/R) (0.5% O_2_; 5% CO_2_; 94.5% N_2_). Cells were treated or, not (DMSO), with 1 mM melatonin (MEL). Rapamycin (RAP) (300 nM) was used to induce of autophagy. 3-methyladenine (3-MA) (5mM) and bafilomycin A1 (BAF A1) (10 nM) were used as inhibitors of phagosome formation and phagosome-lysosome merger, respectively. Cells were treated with the indicated concentrations of MEL or RAP 24 h. (b) Cell viability was measured using the MTT assay and demonstrated as % of control (DMSO normoxia) (c) and the release of β-hCG in the cell media was measured by ELISA and demonstrated as % of control (DMSO normoxia). Data shown are mean ± SD and were analyzed using Student’s *t*-test (* *p*<0.05 and ** *p*<0.01, compared to the same treatment in normoxic conditions; whilst ^#^ *p*<0.05 indicates comparison to vehicle control, DMSO), n=7.

To further investigate the mechanisms involved in the cell viability disruption caused by melatonin in challenging conditions such as H/R, we further investigated the level of phosphorylation of the enzyme AMPKα (Thr 172). The phosphorylation of AMPKα is mainly induced by lower ATP levels [28] or, in cases of hypoxia, by ROS [15]. The phosphorylation of AMPKα was normalised to the total levels of AMPKα, which were steady, irrespective of conditions and treatments. In BeWo cells, phospho-AMPKα levels were increased 6.7 fold in cells exposed to 1 mM melatonin in normoxia, compared to vehicle controls. In BeWo cells exposed to H/R without melatonin, phospho-AMPKα levels were increased 12-fold, compared to vehicle controls. The maximal activation of AMPKα was achieved by melatonin under H/R conditions, where phospho-AMPKα levels were increased 16-fold, compared to all other conditions (**Fig. 2a**). In addition, we investigated the activation of the factor PP2Ac, which has a protective role against hypoxia [29]. The activation of autophagy is among the protective actions driven by the dephosphorylation at Tyr 307 of PP2Ac, which is considered a marker of poor prognosis in malignant cells [13]. Interestingly, BeWo cells show lower levels of phosphorylated PP2Ac when exposed to H/R, independently of melatonin treatment (**Fig. 2b**), compared to normoxia. Beclin-1 is considered a molecular scaffold of autophagy modulators (including AMPKα and PP2Ac) and controls autophagosome formation. Beclin-1 protein levels were increased in BeWo cells exposed to H/R, compared to normoxia. Melatonin did not modulate Beclin-1 levels in either normoxia or H/R conditions. As expected, H/R upregulates upstream regulators of autophagy, with autophagy being inhibited by melatonin in BeWo cells.

**Fig. 2:**
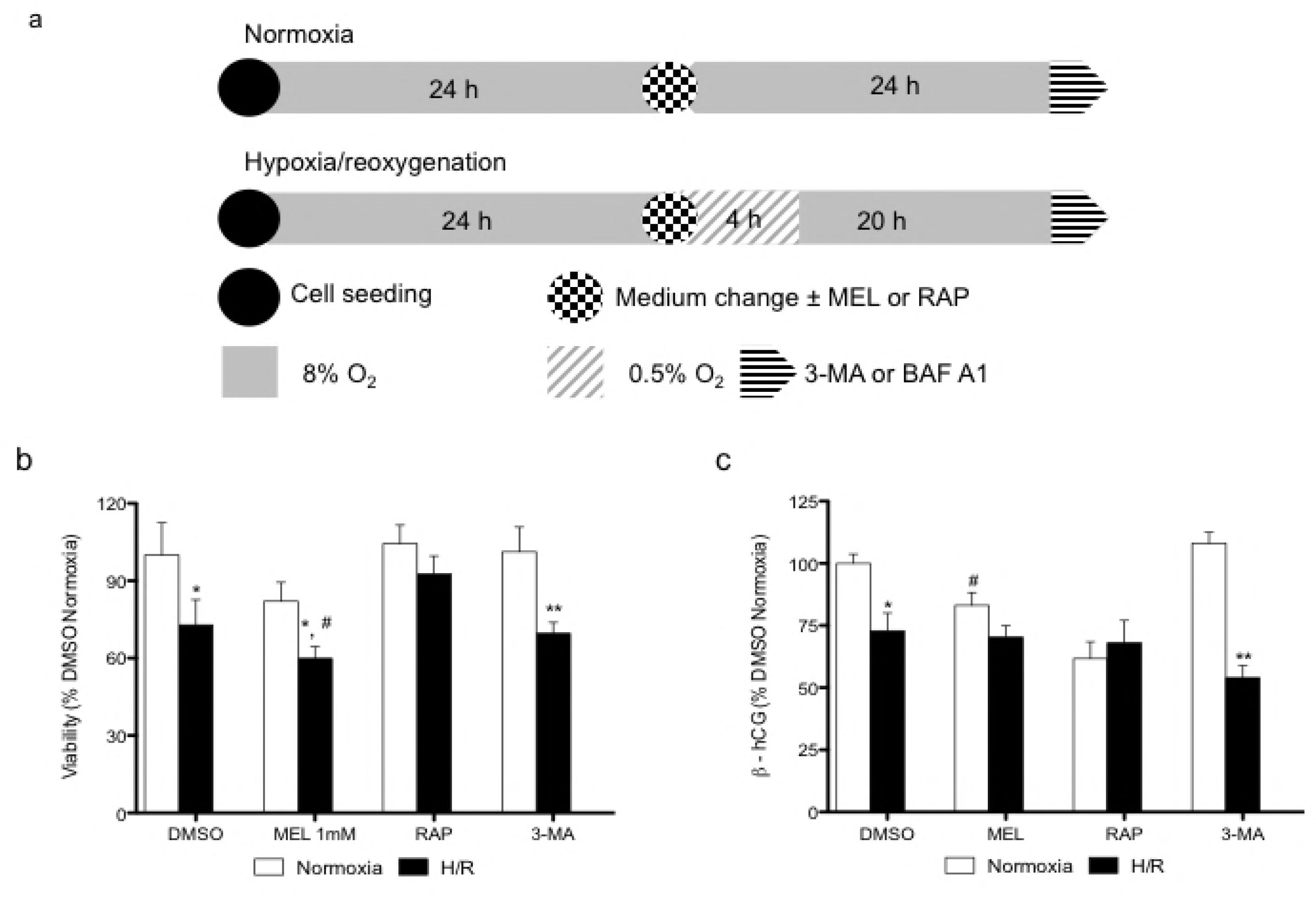
Hypoxia/reoxygenation (H/R) and melatonin modulate upstream autophagic factors in BeWo cells. (a) Relative optical density of phospho-AMPKα (Thr 172), normalized with total AMPKα. The relative molecular mass (kDa) is indicated to the right of the representative blot. (b) Relative optical density of phospho-PP2Ac (Tyr 307), normalized with total PP2Ac. (c) Relative optical density of Beclin-1 with total protein. Total protein amounts were measured by staining with MemCode reversible protein stain. Data are shown as mean ± SD and were analyzed using ANOVA, followed by Student-Newman-Keuls (\**p*<0.05; \*\**p*<0.01; \*\*\**p*<0.001), n = 5.

### Melatonin regulates autophagy in BeWo cells

To determine the downstream pathway involved in the elongation of the autophagosome, we investigated the protein levels of ATG5-ATG12, ATG 16β, ATG 16α, ATG7, and the lipidation of LC3B. The ATG5-ATG12 complex is responsible for the activation (*i.e.* lipidation) of LC3B [30], which is anchored in the autophagosome complex by ATG16 [31]. Rapamycin, which was used as a positive control of autophagy, as expected, increased ATG5-ATG12, ATG16, LC3B-II, and ATG7 under normoxia and H/R conditions, compared to vehicle controls. As with rapamycin, both ATG16 and ATG5-ATG12 protein levels were increased by H/R. Melatonin had no effect on ATG5-ATG12 protein levels, whereas it increased ATG16 in H/R (**Fig. 3a**). ATG7 controls the activation of the whole formation of the autophagosome [2]. ATG7 was significantly increased (by 1.97-fold) in BeWo cells exposed to H/R, in comparison to normoxia. Melatonin increased the ATG7 content by 1.56-fold under normoxia. In H/R conditions, melatonin led to a significant 1.24-fold increase in ATG7 levels, compared to BeWo exposed to DMSO (**Fig. 3a and b**). The activation of LC3B into LC3B-II was significantly increased in BeWo cells exposed to H/R compared to normoxia, with melatonin having no significant effect (**Fig. 3a and 3c**). The lipidation of LC3B is visually characterized by the formation of different puncta, compared to non-active LC3B, which presents with a more uniform cytosolic distribution. As showed in the **figure 3d,** rapamycin and H/R activate the formation of LC3b-puncta, which may indicate the presence of an active autophagosome. Melatonin is not able to significantly activate the whole machinery responsible for the autophagosome formation.

**Fig. 3:**
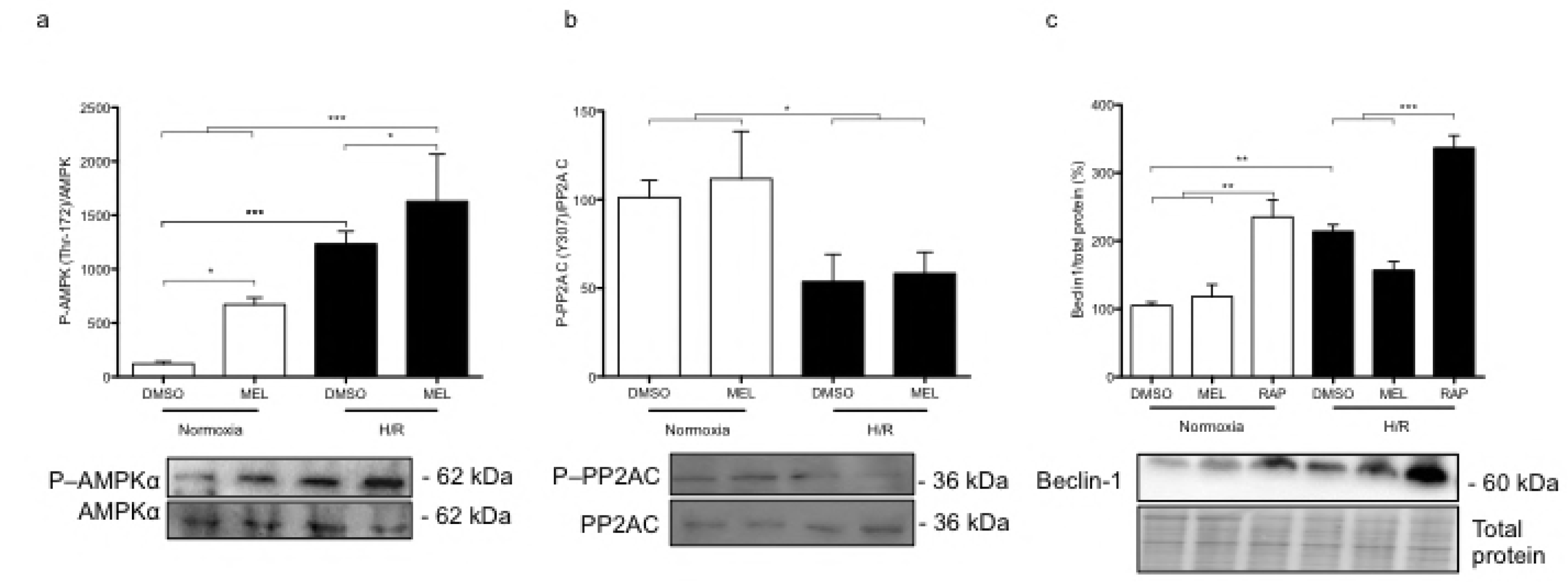
Hypoxia/reoxygenation and melatonin differentially regulate autophagosome formation in BeWo cells. (a) BeWo cells were exposed to normoxia (8% O_2_; 5% CO_2_; 87% N_2_) or hypoxia (4 h) /reoxygenation (20 h) (H/R) (0.5% O_2_; 5% CO_2_; 94.5% N_2_) with or without treatment with melatonin (1 mM) or rapamycin (300 nM). Protein levels of autophagy-related (ATG) ATG5-ATG12, ATG16β, ATG16α, and ATG7 were detected by immunoblotting using monoclonal antibodies. Microtubule-associated proteins 1A/1B lightchain 3B (LC3B) was detected by immunoblotting using a polyclonal antibody. The relative molecular mass (kDa) is indicated to the right of the representative blot. Total protein amounts were measured by staining with MemCode reversible protein stain. ATG16β and ATG16α were represented with two distinct blot images, separated by a dotted line. (b) Relative optical density of ATG7 normalized with total protein is expressed in comparison to vehicle control (DMSO) in normoxia. (c) Relative optical density of lipidated LC3B (LC3B-II) were normalized with total protein being expressed in comparison to vehicle control (DMSO) in normoxia. Data shown are mean ± SD and were analyzed using ANOVA, followed by Student-Newman-Keuls (\**p*<0.05; \*\**p*<0.01; \*\*\**p*<0.001 compared to the same treatment in normoxic conditions, ^#^ *p*<0.05: indicates significance in comparison to DMSO in H/R), n = 5. (d) The cells were fixed and probed for LC3B using immunofluorescence staining, n=4. BeWo cells cultured under normoxia or H/R treated with dimethyl sulfoxide (DMSO) 0.1%, melatonin 1mM or rapamycin 300 nM were fixed in methanol, stained for LC3B (red), counterstained with 4’,6-diamidino-2-phenylindole (DAPI) (nucleus staining blue), and observed by confocal microscopy at 20x magnification.). Scale bar represents 10 μm.

### Melatonin reduces autophagic activity and activation of Nrf2 in BeWo cells exposed to H/R

To further investigate the role of melatonin in the regulation of the autophagy pathway, bafilomycin A1 (BAF A1) was added to the BeWo cell medium during the last 2 h of cell culture under H/R. BAF A1 prevents the fusion of the autophagosome with the lysosome, thereby allowing the investigation of autophagic activity by measuring the protein levels of SQSTM1/P62, which acts as an adaptor for protein trafficking, aggregation, and degradation [32]. **Figure 4a** indicates that, in H/R conditions, BAF A1 increased SQSTM1/P62 by 6.3 fold, compared to BeWo cells under H/R without BAF A1 treatment, suggesting a high level of autophagic activity. In contrast, the BeWo cells exposed to H/R that were treated with melatonin and BAF A1 showed significantly lower levels of SQSTM1/P62, vs cells treated with vehicle control (DMSO), indicating lower autophagic activity. The dashed line represents the protein levels of SQSTM1/P62 in BeWo cells exposed to H/R and treated with rapamycin, as a positive control of the degradation of the autophagic cargo, including SQSTM1/P62. SQSTM1/P62 shares a positive feedback loop with the transcription factor Nrf2 [5], with Nrf2 showing significantly higher protein levels in BeWo cells exposed to H/R compared to normoxia, with Nrf2 levels similar to that of cells exposed to the positive control, sulforaphane, an Nrf2 activator (**Fig. 4b**). BeWo cells exposed to H/R and treated with melatonin, showed significantly less Nrf2 protein levels compared to DMSO, suggesting that melatonin downregulates the Nrf2 protective pathway in these cells. We therefore investigated the Nrf2 nuclear content, which represents the active form of this transcription factor, and found similar results: cells exposed to H/R plus melatonin showed 3-fold reduction in nuclear Nrf2, compared to cells exposed to H/R without melatonin treatment (**Fig. 4d**). As previously described [21], melatonin induces apoptosis in BeWo cells (**Fig. 4c**). The cleavage of poly (ADP-ribose) polymerase (PARP), a hallmark of apoptosis, was increased in BeWo cells exposed to H/R, compared to normoxia. Moreover melatonin in normoxia increased the cleavage of PARP in comparison to the control vehicle. In contrast, cells exposed to the positive control of autophagy, rapamycin, in both normoxia and H/R showed decreased levels of cleaved PARP, suggesting that the induction of autophagy has a beneficial effect on BeWo cell survival, and that the downregulation of autophagy by melatonin contributes to BeWo cell death.

**Fig. 4:**
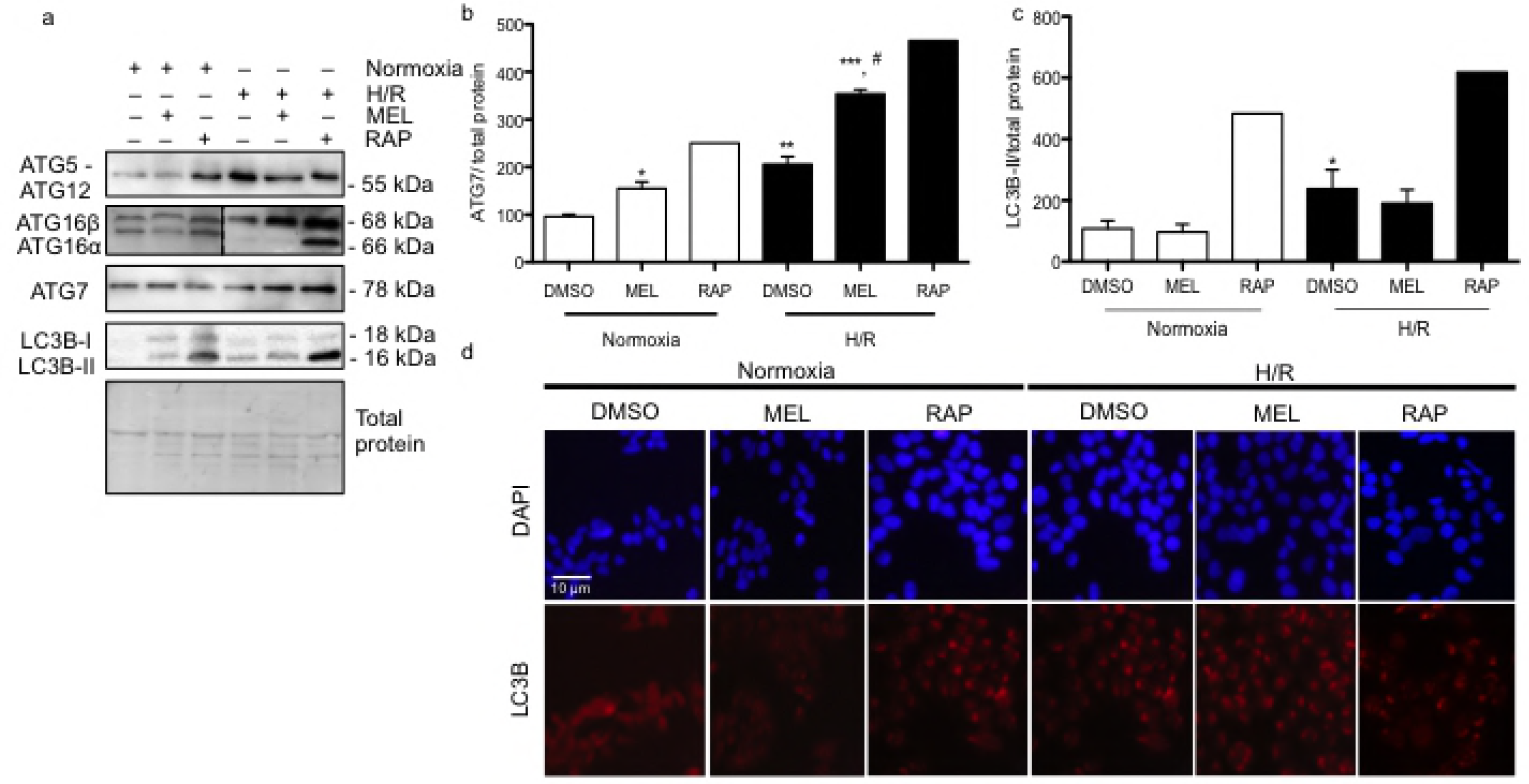
Melatonin reduces autophagy turnover thereby increasing BeWo cell vulnerability to H/R. BeWo cells were exposed to normoxia (8% O_2_; 5% CO_2_; 87% N_2_) or hypoxia (4 h)/reoxygenation (20 h) (H/R) (0.5% O_2_; 5% CO_2_; 94.5% N_2_). (a) Sequestosome-1, SQSTM1/P62, protein expression was detected by immunoblotting using a polyclonal antibody in BeWo cells exposed to H/R. Cells were treated with dimethyl sulfoxide (DMSO) 0.1% or 1 mM melatonin (MEL) for 24 h. Cells were also treated with BAF A1 (10 nM) during the last 2 h of BeWo cell culture. (b) Nrf2 protein expression was detected by immunoblotting using a polyclonal antibody in BeWo cells exposed to normoxia or H/R. Sulforaphane 10 μM was added to BeWo cells for 24 h as an Nrf2 inducer. (c) Relative optical density of the carboxy-terminal catalytic domain of the poly (ADP-ribose) polymerase (PARP), which was cleaved between the Asp214 and Gly215 was normalized with total PARP. Both cleaved PARP and PARP1 were detected by immunoblotting using a polyclonal antibody. (d) The nuclear and the cytosolic contents of the Nrf2 protein of BeWo cells exposed to H/R were detected by immunoblotting. Histone H3 and α-tubulin were used as protein markers of the purity of nuclear and cytosolic fractions, respectively. Representative blots are shown below the graphs and the relative molecular mass (kDa) is indicated to the right of the blot. Total protein amounts were measured by staining with MemCode reversible protein stain. Data shown are mean ±SD and were analyzed using ANOVA, followed by Student-Newman-Keuls (a, b, c) or Student’s *t*-test (d) (\**p*<0.05; \*\**p*<0.01; \*\*\**p*<0.001), n = 4-5.

### Melatonin and hypoxia/reoxygenation decrease the mitochondrial content in BeWo cells

Previous work indicates that melatonin reduces the mitochondrial membrane potential in BeWo cells [21], with mitochondrial content being reduced under hypoxic conditions [33]. In normoxia conditions, melatonin led to a 1.8-fold decrease in the mitochondrial marker TOMM20. As demonstrated in the **figure 5a**, cells exposed to H/R show a 2.5-fold decrease in TOMM20 protein level, compared to normoxia. In H/R conditions, melatonin led to a 4.5-fold decrease in TOMM20, which was significantly lower compared to all other conditions. BAF A1 was added in H/R to block the merging of the autophagosome with lysosome and the consequent catabolism of damaged cellular components. BAF A1 was therefore used as a control to investigate if mitophagy is involved in the reduction of the TOMM20 mitochondrial content under H/R conditions. As demonstrated in the **figures 5a and 5b**, BeWo cells exposed to H/R and treated with BAF A1 presented similar lower mitochondria content in comparison to the normoxia control, suggesting that autophagy was responsible for the reduction of the mitochondrial content observed under H/R. **Figure 5b** shows the colocalization of the mitochondrial marker COX IV and LC3B in BeWo cells. In H/R, cells exposed to melatonin or BAF A1 presented a similar pattern of colocalization of COX IV and LC3B. This may indicate that the decreased mitochondrial content is due to mitophagy in BeWo cells exposed to H/R and treated with melatonin, vs H/R exposure without melatonin.

**Fig. 5:**
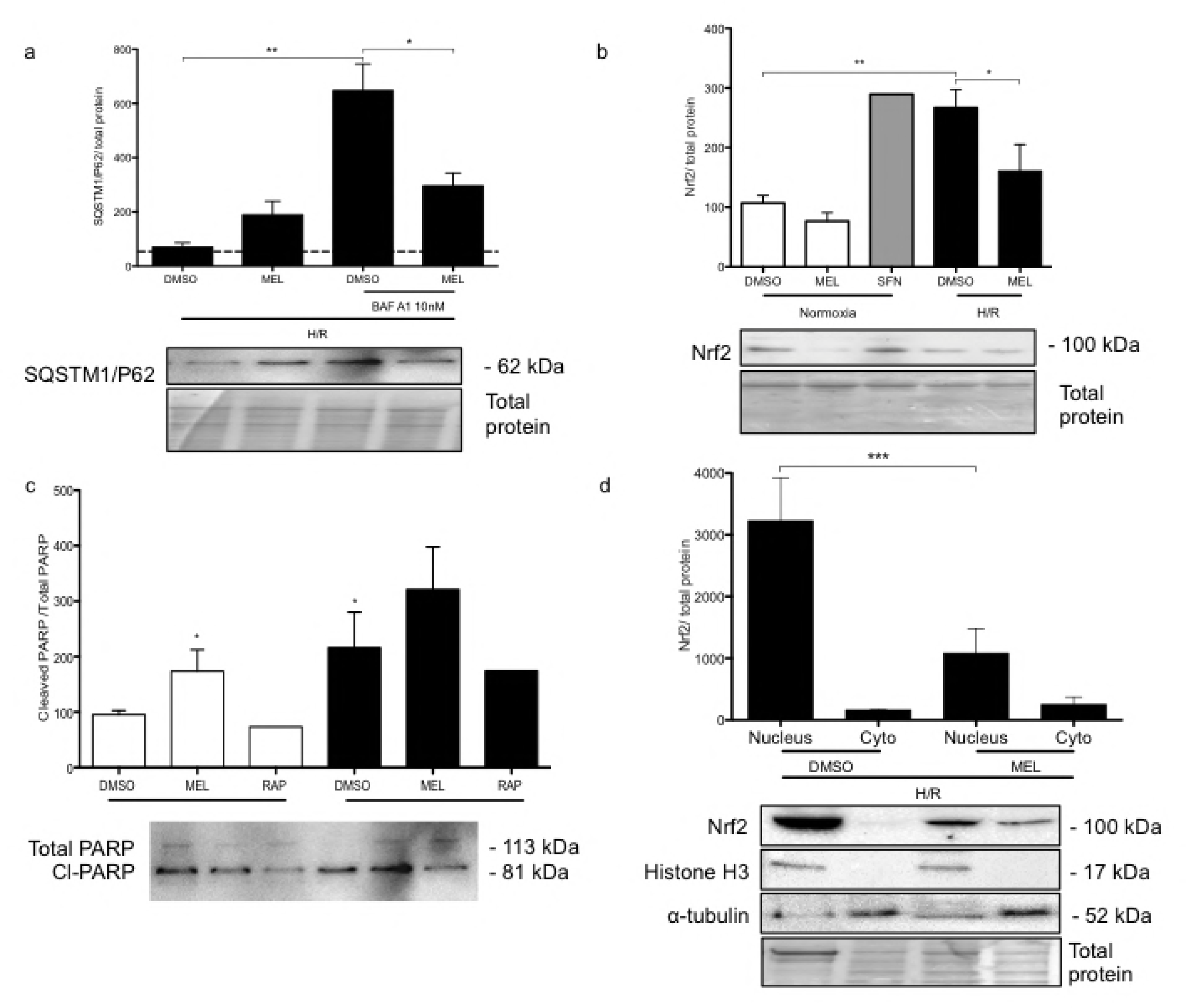
Melatonin and hypoxia/reoxygenation decrease the mitochondrial content in BeWo cells. BeWo cells were exposed to normoxia (8% O_2_; 5% CO_2_; 87% N_2_) or hypoxia (4 h)/reoxygenation (20 h) (H/R) (0.5% O_2_; 5% CO_2_; 94.5% N_2_) with or without treatment with melatonin (1 mM) or bafilomycin (BAF A1: 10 nM). (a) TOMM20 protein level was detected by immunoblotting using a polyclonal antibody. The relative molecular mass (kDa) is indicated to the right of the representative blot. Total protein amounts were measured by staining with MemCode reversible protein stain. Data shown are mean ± SD and were analyzed using ANOVA, followed by Student-Newman-Keuls (\**p*<0.05, compared to DMSO in normoxia; ^#^*p*<0.05, when compared to all other groups), n=4. (b) BeWo cells cultured in H/R conditions and were fixed and probed for cytochrome c oxidase IV (COX IV) and microtubule-associated proteins 1A/1B light chain 3B (LC3B) using immunofluorescence staining. Representative pictures are shown for both COX IV + LC3B staining, COX IV + DAPI staining, and LC3B staining + DAPI. Scale bar represents 10 μm.

### In H/R, melatonin upregulates autophagy in primary vCTBs

As shown in the **figure 6a**, primary vCTBs exposed to H/R with or without treatment with melatonin, had raised protein level of Beclin-1 and LC3B lipidation, compared to normoxia. Intriguingly, in H/R, melatonin treatment increases dramatically the LC3B lipidation (**Fig. 6a, d**). In H/R, primary vCTBs treated with BAF A1, vs not, showed increased SQSTM1/P62 protein levels, suggesting that the autophagic turnover was activated. The utilization of BAF A1 showed melatonin to have an additive effect on autophagic turnover (**Fig. 6b**). Nrf2 was also significantly increased (by 2.25-fold) in primary vCTBs exposed to H/R, compared to normoxia (**Fig. 6a, c, d**). Following melatonin treatment, Nrf2 expression was further upregulated (by 1.44-fold), vs H/R only exposure. Moreover, melatonin led to the nuclear translocation of Nrf2 and to the lipidation of LC3B, as shown in the **figure 6d**. Both the increased lipidation of LC3B and the nuclear translocation of Nrf2 indicate a protective role of melatonin in human vCTB under H/R.

**Fig. 6:**
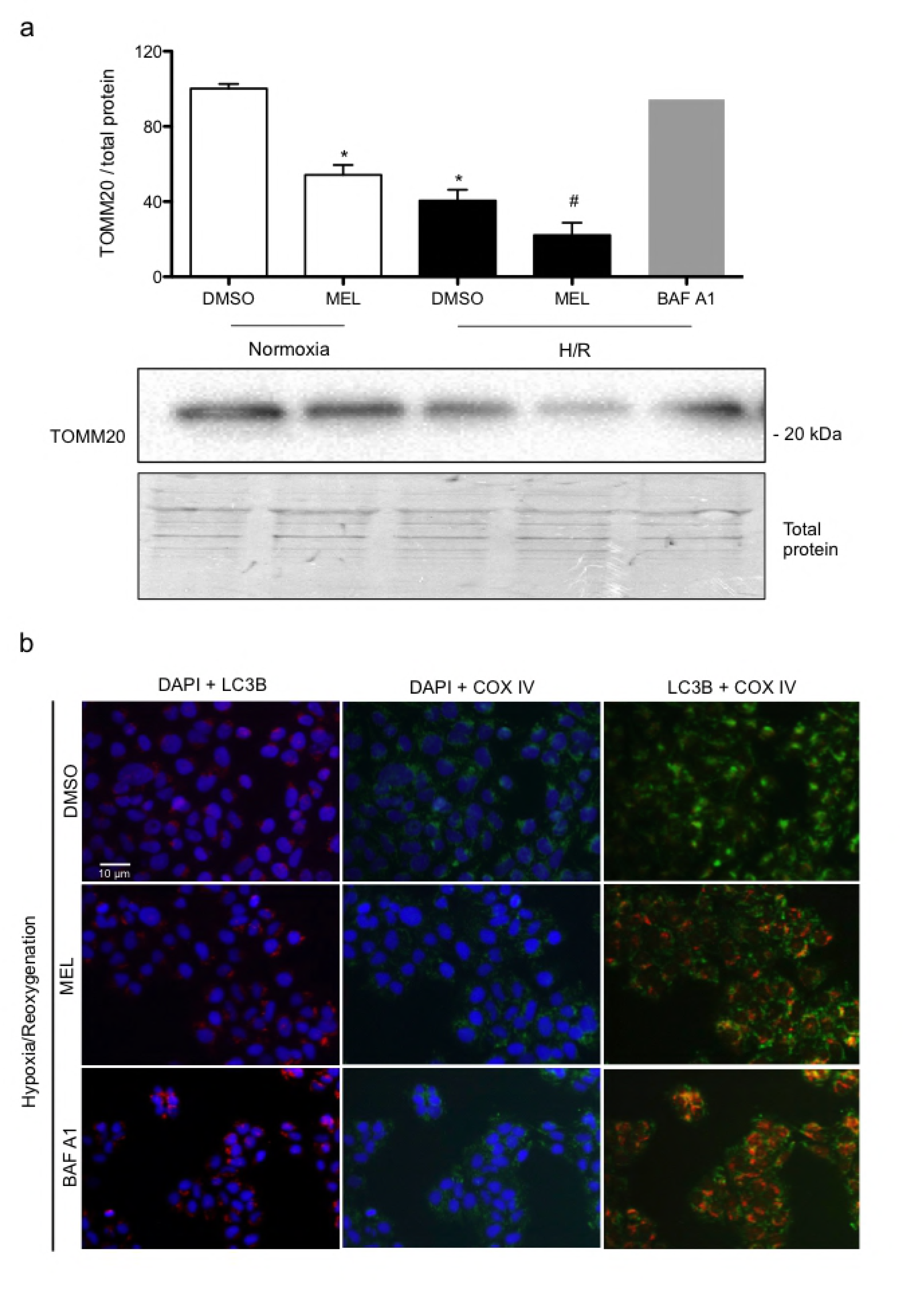
Melatonin increases autophagy turnover and Nrf2 expression in primary villous cytotrophoblasts exposed to hypoxia/reoxygenation. Human primary villous cytotrophoblasts (vCTBs) were exposed to normoxia (8% O_2_; 5% CO_2_; 87% N_2_) or hypoxia (4 h) /reoxygenation (20 h) (H/R) (0.5% O_2_; 5% CO_2_; 94.5% N_2_). Cells were treated with dimethyl sulfoxide (DMSO) 0.1% or 1 mM melatonin (MEL) for 24 h. (a) Protein expression of nuclear factor (erythroid-derived 2)-like 2 (Nrf2), Beclin-1, and microtubule-associated proteins 1A/1B lightchain 3B (LC3B) were detected by immunoblotting. (b) Sequestosome-1 is also known as the ubiquitin-binding protein p62 (SQSTM1/P62), and its protein expression was detected by immunoblotting using a polyclonal antibody in vCTB exposed to H/R. Cells were treated with dimethyl sulfoxide (DMSO) 0.1% or 1 mM melatonin (MEL) for 24 h. vCTB were also treated with bafilomycin A1 (10 nM) during the last 2 h of culture. Total protein amounts were measured by staining with MemCode reversible protein stain. The relative molecular mass (kDa) is indicated to the right of the representative blot. (c) Relative optical density of Nrf2 normalized with total protein. Data shown are mean ± SD and were analyzed using ANOVA, followed by Student-Newman-Keuls (\**p*<0.05; \*\*\**p*<0.001), n = 4. (d) vCTB cells were fixed and probed for Nrf2 (green) and LC3B (red) using immunofluorescence staining. 4’,6-diamidino-2-phenylindole (DAPI; blue) was applied as a nucleus stain. Scale bar represents 10 μm.

## Discussion

H/R disturbs the cellular homeostasis; leading to the activation of survival mechanisms, such as autophagy and the transcription factor Nrf2 [6, 34]. Most data indicates melatonin to be protective to normal cells and cytotoxic to cancer cells [19, 35]. In accord with these studies, the present study shows that H/R is able to activate both autophagy and Nrf2 in normal and tumoral trophoblasts, with melatonin inhibiting autophagy and Nrf2 in the BeWo choriocarcinoma cell line BeWo, whilst upregulating these pathways in human primary vCTBs. The above results demonstrate, for the first time, that autophagy is activated by H/R, but inhibited when treated with concomitant melatonin treatment in BeWo cells. Despite an increased total and nuclear protein fraction in BeWo cells exposed to H/R, Nrf2 activation is inhibited by melatonin resulting in the activation of cell death via the cleavage of PARP-1. In contrast, human primary vCTBs exposed to H/R and treated with melatonin showed increased autophagy and Nrf2 activity, which corroborates previous data showing a protective role of melatonin for normal placental cells [20, 36].

Previous work has shown melatonin to be produced, and to have important protective functions, in primary human trophoblasts as well as in the pregnancy [20, 21, 36, 37]. H/R increases oxidative stress levels and lowers primary villous trophoblast viability, with melatonin affording protection against such H/R effects ([20]; Fagundes, et al. *in press*). In the current study, BeWo cells exposed to H/R showed reduced cell viability and lower hCG release (a marker of the endocrine function of normal and tumoral placental trophoblasts), compared to normoxia [38]. The treatments with melatonin or 3-MA, an inhibitor of autophagy, were not able to rescue the levels of viability or hCG, whereas rapamycin, which is largely used as an activator of autophagy via mTOR inhibition, rescued the viability of BeWo cells exposed to H/R. Importantly, the current results show that the upstream modulators of autophagy, PP2Ac and AMPKα, are activated under H/R conditions. Such data on cell viability and rapamycin effects implicate autophagy in the survival of BeWo choriocarcinoma cells challenged by H/R. Intriguingly, the activation of LC3B in its lipidated form, LC3B-II, was not significantly modulated by H/R in BeWo cells. This could be due to the high basal activity of autophagy already described in cancer cell models exposed to hypoxic conditions, where results indicate increased activation of LC3B only after longer hypoxia durations (e.g. 48 h of 0.1% O_2_) [39].

Melatonin induces BeWo cell death, in contrast to the effects of autophagy activation by rapamycin. This could suggest that melatonin inhibits autophagy in BeWo cells, which is supported by the present results showing decreased SQSTM1/P62 protein level in BeWo cells treated with melatonin or BAF A1. Coto-Montes et al. [40] proposed that melatonin inhibits autophagy via direct inhibition of upstream factors, such as the inflammatory factor, c-Jun N-terminal kinase (JNK). Consequent to JNK inhibition, autophagy activation has been shown to be decreased in a hepatoma cell line [41]. Other studies indicate that the inhibitory effect of melatonin on autophagy may be mediated by the activation of endoplasmic reticulum (ER)-stress [42, 43]. The lack of variation in the levels of LC3B lipidation in melatonin-treated BeWo cells is not related to the lack of autophagic activity, but rather arises as a consequence of the active destruction of defective mitochondria via mitophagy [39]. We observe a reduction in mitochondrial content in melatonin treated BeWo cells exposed to H/R, which may indicate heightened levels of mitophagy, and which has been shown to be harmful for cancer cells [44, 45]. Our result showing the decreased mitochondrial content of cancer cells exposed to H/R and treated with melatonin corroborates other studies that have found antiproliferative and pro-apoptotic activity of melatonin, evident in both *in vitro* and *in vivo* models as well as in clinical assays for an array of diverse cancers [19, 42, 43].

In both normal and cancer cells, Nrf2 activation induces the expression of antioxidant enzymes [6]. In H/R conditions, melatonin reduces the nuclear content of Nrf2 in BeWo cells, thereby lowering the cells levels of protective antioxidants and contributing to increased cleavage of PARP and cell death. In contrast, vCTBs exposed to H/R and treated with melatonin showed increased nuclear translocation of Nrf2. As such, the differential effects of melatonin in normal and tumoral trophoblasts is associated with parallel changes in Nrf2 and autophagy, with both being decreased in H/R-exposed BeWo cells, but increased in H/R-exposed vCTB. The dual role of melatonin in such important mechanisms of cell defense – Nrf2, and autophagy – indicates a co-ordinated set of changes that is harmful in tumoral cells and protective in normal cells. The differential effects of melatonin in tumoral vs normal cells have been shown over a number of decades, leading to great interest in the anticancerous utility of melatonin, especially given its low toxicity, small size, and amphiphilic profile [17, 18]. However, the mechanisms underlying the differential effects of melatonin in normal vs cancer cells is still poorly understood. In breast cancer cells, melatonin alters the patterned expression of microRNAs and as well as DNA methylation. It requires investigation as to whether melatonin has significant differential impacts on such epigenetic processes in normal vs tumoral cells, including placental cells [35, 46]. Recent work by Hardeland highlights the importance of melatonin acts as an epigenetic modulator in different cell types [35], although this has still to be investigated in placental cells (**Figure 7**). In a recent review by Reiter et al. [19], the cytotoxicity of melatonin in cancer cells was described as possibly via receptor-dependent and receptor-independent actions, with both mechanisms leading to decreased cancer cells viability by inhibiting protective pathways that would be activated in suboptimal conditions, such as hypoxia and low levels of amino acids.

**Figure 7:**
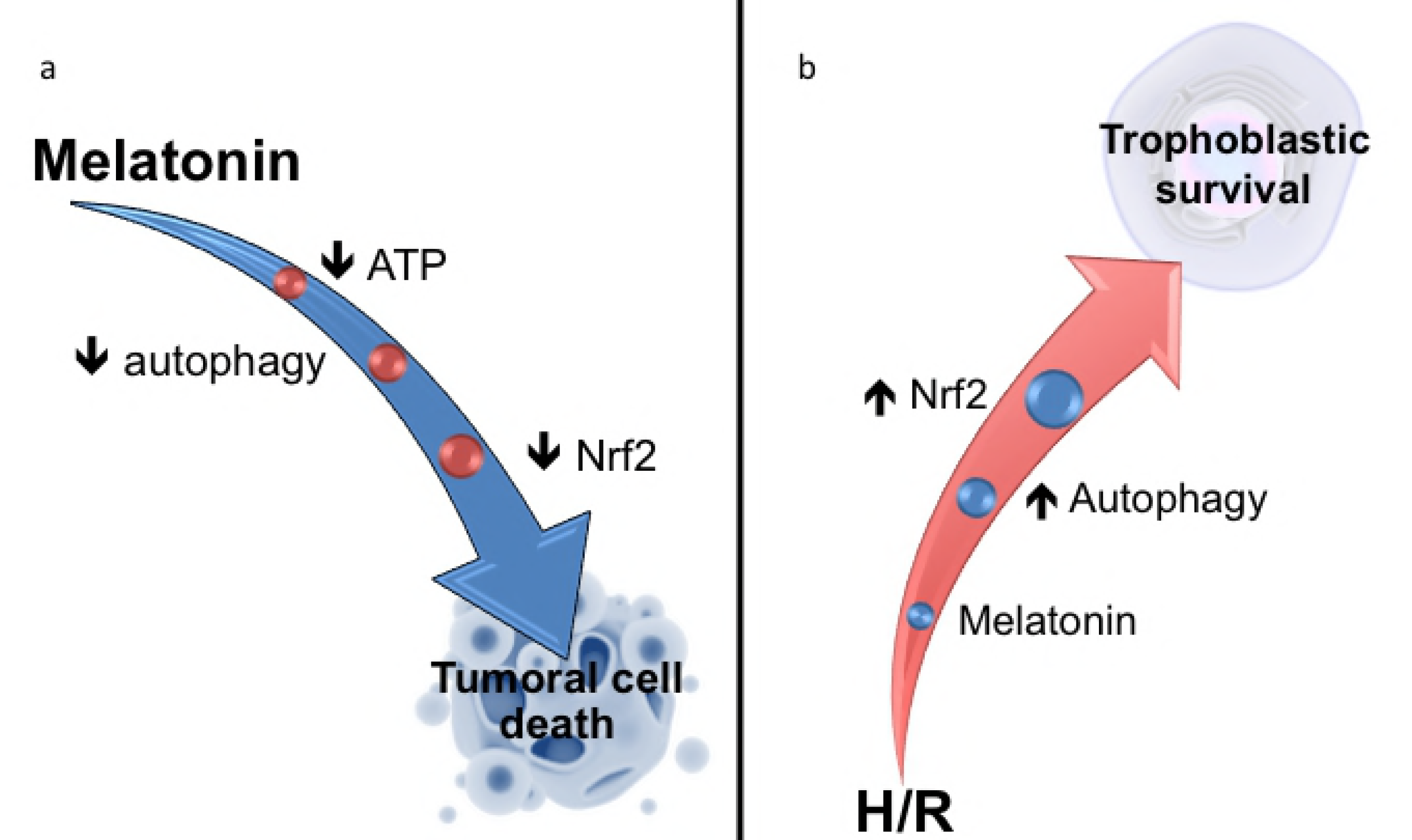
Melatonin acts differentially in tumoral and normal placental cells. (a) In the BeWo choriocarcinoma cell line, melatonin increases the phosphorylation of 5 ′adenosine monophosphate-activated protein kinase (AMPK), due to the reduction of ATP levels [<u>50</u>]. Melatonin inhibits autophagic activity as well as the activation of the nuclear factor (erythroid-derived 2) -like 2 (Nrf2) transcription factor, contributing to BeWo cell death. (b) In primary cell cultures of human placental villous cytotrophobasts (vCTBs), H/R disrupts the cellular homeostasis. Melatonin treatment induces activation of two pro-survival mechanisms: autophagy and Nrf2, which help to restore homeostasis and cell survival.

This study makes an important contribution to the understanding of the biological underpinnings that result in the differential effects of melatonin in tumoral vs normal placental cells. The data indicate that the differential effects of melatonin on Nrf2 and autophagy are intimately linked to this. Future research should investigate as to whether the changes in Nrf2 and autophagy are linked to epigenetic processes (such as microRNAs), mitochondrial factors (such as oxidative phosphorylation vs glycolysis) and alterations in endogenous melatonergic pathway regulation. In conclusion, the data show that the differential effects of melatonin on Nrf2 and autophagy are important to its protection of normal cells and increased apoptosis of tumor cells, including under H/R conditions.

## Acknowledgments

The authors thank the women who donated their placentas for this study.

## Supporting information

**S1 Table**. Antibodies used for Western blot analyses

**S2:** Raw data of all of the results presented in this manuscript.

Author contributions
All co-authors contributed to the design of the study protocol. LSF and JBP were involved in data acquisition. LSF, was actively involved in the writing of the manuscript assisted by CV. All co-authors provided critical revisions and their final approval for the manuscript’s publication in PLOS ONE. This manuscript is not under review elsewhere and all authors are in agreement with the contents of this manuscript.

## Notes

**Funding** Supported by grants from: the Natural Sciences and Engineering Research Council of Canada (NSERC) (no. 262011) to CV; NSERC: Undergraduate Student Research Award (URSA) studentship awards to JBP; *the Ministère de l’Éducation, du loisir et du sport* (MELS) *du Québec-Fonds de recherché du Québec-Nature e technologies* (FRQNT) and also studentship from *Fondation Armand Frappier to LSF*; and from *Fondation Armand Frappier* and *Réseau Québécois en reproduction* (RQR)-NSERC-Collaborative Research.

